# Validation of a habitat suitability index for oyster restoration

**DOI:** 10.1101/039875

**Authors:** Seth J. Theuerkauf, Romuald N. Lipcius

## Abstract

Habitat suitability index (HSI) models provide spatially explicit information on the capacity of a given habitat to support a species of interest, and their prevalence has increased dramatically in recent years. Despite caution that the reliability of HSIs must be validated using independent, quantitative data, most HSIs intended to inform terrestrial and marine species management remain unvalidated. Furthermore, of the eight HSI models developed for eastern oyster (*Crassostrea virginica*) restoration and fishery production, none has been validated. Consequently, we developed, calibrated, and validated an HSI for the eastern oyster to identify optimal habitat for restoration in a tributary of Chesapeake Bay, the Great Wicomico River (GWR). The GWR harbors an unparalleled, restored oyster population, and therefore serves as an excellent model system for assessing the validity of the HSI. The HSI was derived from GIS layers of bottom type, salinity, and water depth (surrogate for dissolved oxygen), and was tested using live adult oyster density data from a survey of high vertical relief reefs (HRR) and low vertical relief reefs (LRR) in the sanctuary network. Live adult oyster density was a statistically significant sigmoid function of the HSI, which validates the HSI as a robust predictor of suitable oyster reef habitat for rehabilitation or restoration. In addition, HRR had on average 103-116 more adults m^-2^ than LRR at a given level of the HSI. For HRR, HSI values ≥0.3 exceeded the accepted restoration target of 50 live adult oysters m^−2^. For LRR, the HSI was generally able to predict live adult oyster densities that meet or exceed the target at HSI values ≥0.3. The HSI indicated that there remain large areas of suitable habitat for restoration in the GWR. This study provides a robust framework for HSI model development and validation, which can be refined and applied to other systems and previously developed HSIs to improve the efficacy of native oyster restoration.

## Introduction

Habitat suitability indices (HSI) are a commonly developed and often robust spatially explicit, decision support model used to identify the capacity of a given habitat to support a species of interest (U. S. Fish and Wildlife Service 1981, Roloff and Kernohan 1999). In 1981, the United States Fish and Wildlife Service proposed and developed the first HSI models, which were intended to quantify the value of habitats when considering management alternatives in species-specific conservation and restoration (U. S. Fish and Wildlife Service 1981). HSIs are commonly generated through application of wildlife-habitat relationships to relevant geospatial environmental data within a Geographic Information System (GIS) to develop a composite HSI score with a range of 0 to 1, representing unsuitable to optimal habitat (Brooks 1997). Depending on the relevance of the selected habitat variables, quality of the geospatial environmental data, and reliability of the applied wildlife-habitat relationships, these models can serve as robust spatial tools to inform species management.

Although the USFWS emphasized the need for validation of HSIs, or the quantitative assessment of an HSI’s ability to predict habitat suitability via an independent data set, most HSIs intended to inform the management of terrestrial and marine species have not been validated (Brooks 1997, Araújo and Guisan 2006). Recent HSI validation studies have indicated that unvalidated HSIs can be unreliable indicators of habitat quality (Reiley et al. 2014). Although widely used, HSI models have been criticized as unreliable and lacking scientific rigor (Cole and Smith 1983, Roloff and Kernohan 1999). Despite the potential cost associated with obtaining these independent datasets, the validation process is required to determine the reliability and utility of these models if they are to inform species conservation and management (Brooks 1997, Tirpak et al. 2009, Reiley et al. 2014).

To implement HSI models confidently, they must be tested for accuracy in a four-step process (Brooks 1997, Tirpak et al. 2009, Reiley et al. 2014): development, calibration, verification, and validation. Development involves the use of wildlife-habitat relationships to generate an HSI ranging from 0, representing unsuitable habitat, to 1, representing optimal habitat. Calibration aims to ensure that the HSI spans the full range of values from 0 to 1. Brooks (1997) notes: “The intent is for sites of excellent habitat quality to receive high scores (e.g., 0.7-1.0), and sites of poor habitat quality to receive low scores (e.g., 0-0.3). If the HSI scores do not ordinate across the entire range of values from 0 to 1, then they will be of little use in describing differences among sites.” Verification entails assessment of performance of an HSI model against independent qualitative or categorical (ranked) data. A positive correlation of ranked data, such as presence/absence, and HSI values would provide verification. Validation involves testing the performance of an HSI model against independent quantitative data in space and time, such as against population density or abundance. If validation has been accomplished, verification is not necessary. A note of caution pertains to the use of different, non-independent data for validation. For example, a dataset for a single ecosystem, such as a tributary, could be split in half. One half of the dataset could be used to develop an HSI, and the second half used to test the fit of the HSI model. This would not constitute validation because the data used to test the HSI are not statistically independent of the data used to develop the model.

Few HSIs developed for marine or estuarine species of conservation interest have been verified or validated through use of independent data. Brown and Hartwick (1988) were the first to validate an HSI for a marine species, though it was for aquaculture of the introduced Pacific oyster *Crasostrea gigas* in British Columbia, Canada, not for native oyster restoration. Brown et al. (2000) verified HSI models for various fish and invertebrate species (alewife *Alosa psuedoharengus,* American sand lance *Ammodytes americanus,* Atlantic salmon *Salmo salar,* Atlantic tomcod *Microgadus tomcod,* common mummichog *Fundulus heteroclitus,* winter flounder *Pleuronectes americanus,* American lobster *Homarus americanus,* and soft-shell clam *Mya arenaria)* in an estuarine system. Validation of HSI models was also accomplished for the distribution of juvenile Atlantic salmon *S. salar* in a river (Guay et al. 2000) and coastal aquaculture farms of Pacific oyster *C. gigas* (Cho et al. 2012). Note that in this study, we focus on HSI models, and not on species distribution models such as MAXENT (Phillips and Dudík 2008), which have also been used to predict the distribution of marine species as a function of habitat features (e.g., Elsäßer et al. (2010)).

Numerous HSI models have been developed to guide aquaculture, fishery production, and restoration of oyster species (Table 1). The first to derive a model of habitat quality for an oyster species was Galtsoff (1964) for the eastern oyster *C. virginica.* Galtsoff (1964) developed a mathematically simple model to evaluate potentially productive oyster bottom in Gulf of Mexico and south Atlantic coastal habitats. He used the sum of bottom condition, water movement, water temperature, salinity, food availability, sedimentation, diseases, competition, predation, and pollution, each of which ranged from 0-10. The model integrated singular point measurements for each variable to generate a composite score that would characterize a broad region and therefore cannot be considered an HSI. Galtsoff (1964) recognized that the model was overly simplistic because it weighted all of the variables equally and because the model needed to be revised for different geographic areas. Unfortunately, the model was additive in its habitat quality elements, such that a value of 0 in one habitat characteristic did not produce an index of 0, unlike contemporary HSI models.

**Table 1.** Habitat suitability index models developed for oyster aquaculture, fishery production, and restoration. Calibration is achieved when the HSI approximately spans the full range from 0 to 1. Verification is achieved when the HSI values are positively correlated with independent qualitative or categorical (ranked) data, such as presence/absence data. Validation is achieved when the HSI values correlate positively with independent quantitative data in space and time, such as population density or abundance. If validation has been accomplished, verification is not necessary.

| Authors | Species | Calibrated? | Verified? | Validated? |
| --- | --- | --- | --- | --- |
| *Aquaculture* |  |  |  |  |
| Brown and Hartwick (1988) | *C. gigas* | No * | Yes † | Yes |
| Cho et al. (2012) | *C. gigas* | No * | Yes † | Yes |
| *Fishery production* |  |  |  |  |
| Cake (1983) | *C. virginica* | Yes | No | No |
| Soniat and Brody (1988) | *C. virginica* | Yes | Yes | No |
| *Restoration* |  |  |  |  |
| Battista (1999) | *C. virginica* | Unknown § | No | No |
| Barnes et al. (2007) | *C. virginica* | Yes | No | No |
| Starke et al. (2011) | *C. virginica* | Yes | No | No |
| Pollack et al. (2012) | *C. virginica* | Unknown § | Yes ‡ | No |
| Soniat et al. (2013) | *C. virginica* | Yes | No | No |
| Swannack et al. (2014) | *C. virginica* | Yes | No | No |
| This study | *C. virginia* | Yes | Yes † | Yes |

\* The HSI was not calibrated because index values did not span the range from 0 to 1.
§ The HSI calibration status is unknown because index values are reported as a range from low to high, and not from 0 to 1.
† The HSI was not verified before being validated, but because it was validated, it is also considered to be verified.
‡ Abundance data was collected using an oyster dredge, which provides qualitative, not quantitative, estimates of density. Consequently, the HSI was verified, not validated.

Cake (1983) developed the first true HSI for the eastern oyster in the Gulf of Mexico, meaning that it was intended to be employed using spatially-explicit datasets to determine relative suitability of locations within a given system for fishery production. The Cake (1983) model represents a comprehensive oyster HSI that incorporates a broad suite of relevant environmental variables (Table 2). Soniat and Brody (1988) utilized spatially-explicit environmental and oyster density datasets collected for 38 reef and non-reef sites to apply and verify the Cake (1983) HSI for Galveston Bay, Texas. Using oyster density data, the authors attempted to validate the original Cake (1983) model using a regression approach. However, the authors encountered the issue of lack of statistical independence due to oyster density (their response variable) also being incorporated as a variable in their model, and therefore recognized that the model could not be considered validated. The authors did benefit from removing oyster density from the model, testing the relationship between oyster density and various combinations of the variables incorporated into the original model (i.e., a model verification procedure), and subsequently developed a modified HSI that incorporated the significant explanatory variables.

**Table 2.**
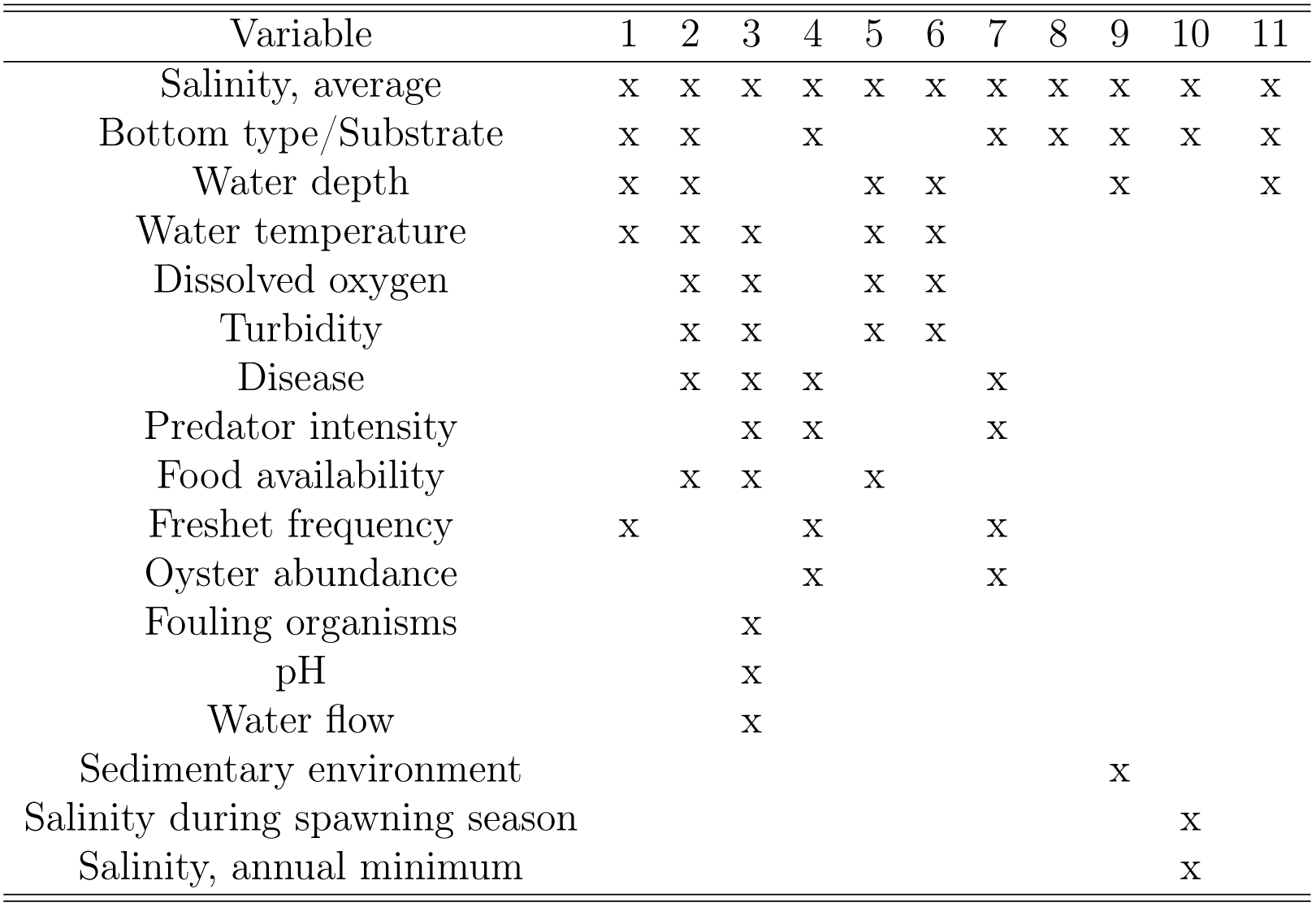
Variables used in habitat suitability index models developed for oyster aquaculture, fishery production and restoration, and the models using each variable. 1-Barnes et al. (2007), 2-Battista (1999), 3-Brown and Hartwick (1988), 4-Cake (1983), 5-Cho et al. (2012), 6-Pollack et al. (2012), 7-Soniat and Brody (1988), 8-Soniat et al. (2013), 9-Starke et al. (2011), 10-Swannack et al. (2014), 11-This study.

With the advent of contemporary geographic information systems (GIS) software, many HSIs for oysters have been developed within the past decade (Table 2). The variables included within these models have varied widely, however, the most common variables utilized are salinity, bottom type or substrate, and water depth. Other variables, such as predator intensity, food availability, or pH are of unknown relevance for all systems and their integration into HSI models requires additional spatially-explicit datasets that rarely exist for most systems. Interestingly, the two most recent published models (Soniat et al. 2013, Swannack et al. 2014) and this study have converged on a simplified model structure that includes varied combinations of the most common variables utilized in previous HSIs (i.e., salinity, bottom type or substrate, and water depth), although the Soniat et al. (2013) and Swannack et al. (2014) models do not include water depth, which is a critical variable for Chesapeake Bay subtidal oyster reefs. Additionally, of the eight published oyster restoration HSIs, none has been validated using an independent, quantitative population dataset. Thus, the reliability of these HSI models for informing oyster restoration and management remains uncertain.

We developed, calibrated and validated an HSI for the eastern oyster in a tributary of Chesapeake Bay, the Great Wicomico River (GWR). In 2004, the U.S. Army Corps of Engineers (ACE) constructed 34.4 ha of sanctuary oyster reef within the system (Schulte et al. 2009). Today, the GWR harbors an unparalleled, restored oyster population (Schulte et al. 2009), and therefore serves as an excellent model system for assessing the validity of an oyster HSI. Here, we describe the development of a simple, reliable HSI for the eastern oyster and present the results from a direct field validation of the model.

## Material and methods

### Study area

The GWR is a tributary on the western shore of the lower Chesapeake Bay (Figure 1). The GWR is located approximately 10 km south of the Potomac River and 25 km north of the Rappahannock River, and has a small watershed consisting predominately of forested and agricultural lands (Southworth et al. 2010). The GWR is mesohaline and is considered a trap-type estuary with gyre-like water circulation patterns that has contributed to its history of significant natural oyster recruitment (Andrews 1979, Southworth et al. 2010). The system is characterized by a single, central deep channel with an extensive sand shoal near the river mouth (Southworth et al. 2010). Oysters within the system exist on public oyster grounds, private lease areas, and no-harvest oyster sanctuaries (Schulte et al. 2009, Southworth et al. 2010).

**Figure 1.**
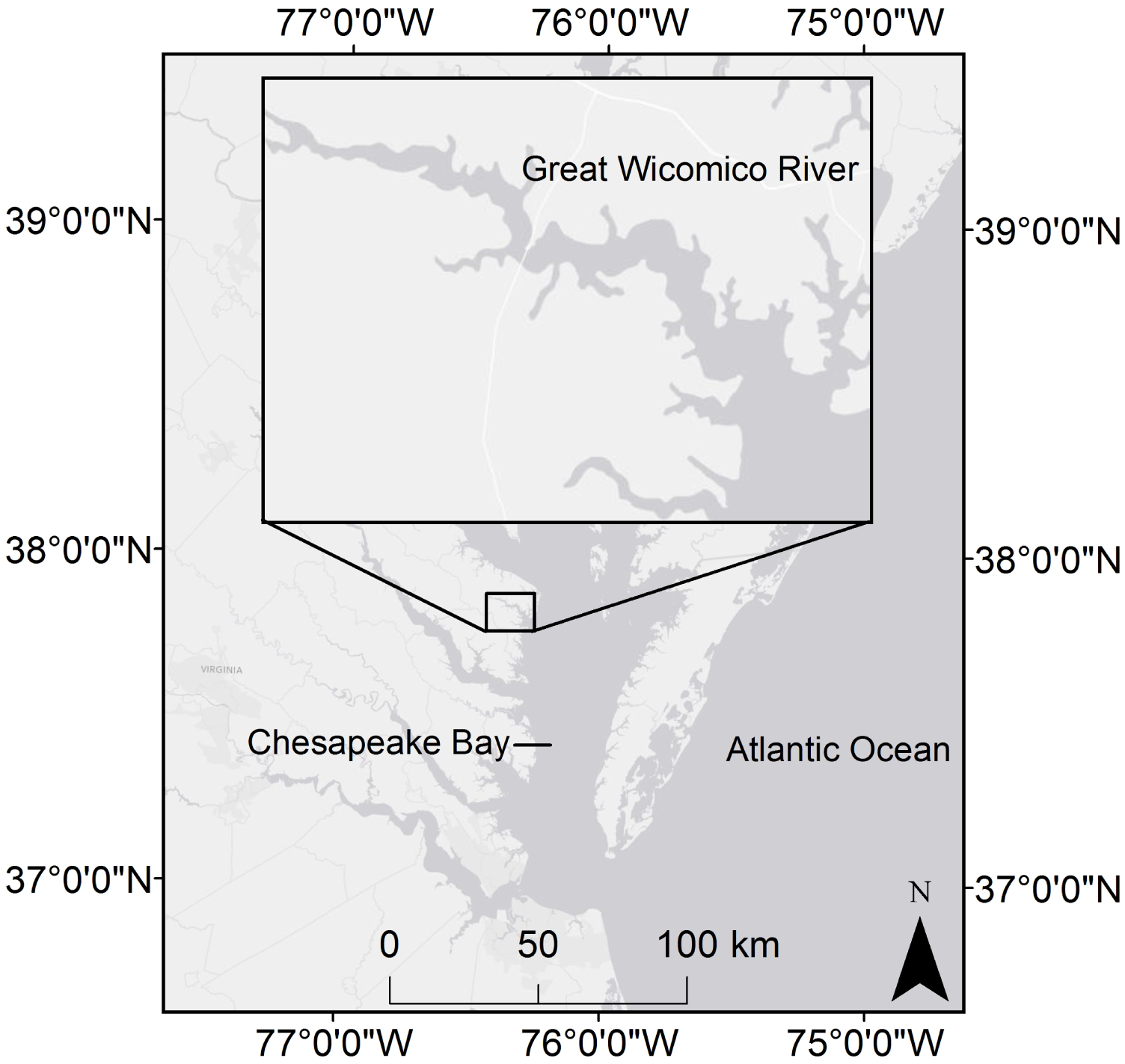
Map showing the location of the Great Wicomico River in relation to Chesapeake Bay.

### HSI development

This HSI was developed in ESRI ArcGIS 10.1 (Environmental Systems Research Institute 2011), and followed a standard logical framework used in the development of previous HSI models (Cake 1983, Battista 1999), except that we added a validation step with independent survey data. The steps in HSI development were as follows:

(i) Assimilation of data sets on environmental variables (e.g., salinity);
(ii) Assessment of habitat requirements for eastern oyster from a literature review;
(iii) Construction of ArcGIS environmental layers;
(iv) Formulation of suitability functions for each of the environmental variables;
(v) Calculation of the HSI for the river system;
(vi) Collection of eastern oyster abundance data for the system from a population survey; and,
(vii) Comparison of HSI values and oyster abundance.

The HSI was derived from Geographic Information System (GIS) layers of environmental and biotic variables, including bottom type, land use, salinity, existence of private oyster leases and public oyster grounds, seagrass cover, dissolved oxygen, and water depth for most of the GWR at depths deeper than 2 m. From these variables, we selected those of greatest relevance to site suitability for oyster restoration in the GWR and for which there was river-wide data, which included bottom type, depth, and salinity (Figure 2). Bottom type was included in this study as oyster reefs constructed on sand, rock or oyster shell are less likely to subside than those built on mud or silt bottom (Schulte et al. 2009). Depth was included as benthic community abundance and diversity decline significantly at low dissolved oxygen (DO) levels (Long and Seitz 2009, Seitz et al. 2009), and depth is a robust surrogate for DO (Powers et al. 2005). During the summer in western shore tributaries of Chesapeake Bay, such as the GWR, DO remains high down to about 4 m (Seitz et al. 2009), after which it declines sharply to 0 mg 1^−1^ around 5 m. This general trend is due to seasonal benthic anoxia and hypoxia events induced by decomposition of plankton and subsequent stratification of the water column (Kemp et al. 2005). Salinity was included as the eastern oyster cannot tolerate extremely low or high salinities for long periods, and prefers upper mesohaline to polyhaline salinities (Battista 1999, Barnes et al. 2007, Carnegie and Burreson 2011, Battista 1999).

**Figure 2.**
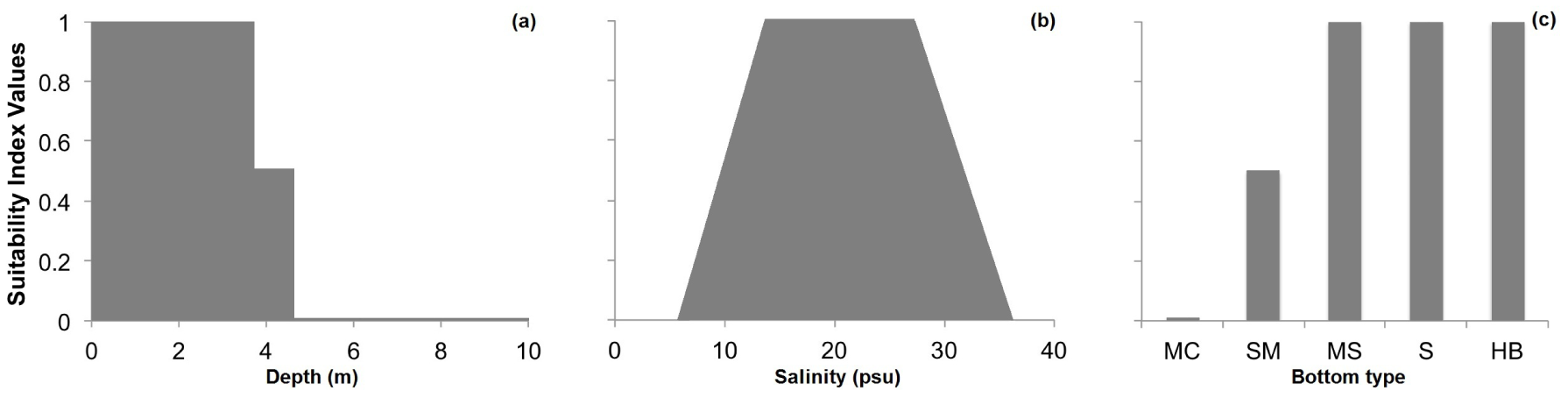
Relationships between the actual value of environmental variables and associated habitat suitability values for (a) water depth (surrogate variable for dissolved oxygen), (b) salinity, and (c) bottom type (MC = muddy clay, SM = sandy mud, MS = muddy sand, S = sand, HB = hard bottom). Scores of 0 are unsuitable and values of 1 are optimal.

Data for GIS layers of bottom type were derived from multiple sources including: 1) a 2009 NOAA acoustic seabed mapping survey (1 m × 1 m resolution), and 2) field notes taken by VIMS researchers during the 2012 reef monitoring sampling of the ACE reefs (Bruce et al. 2010). Bottom type data derived from the 2009 NOAA acoustic seabed mapping survey included the majority of downriver bottom, but excluded inshore areas shallower than 2 m due to the inability of the research vessel to navigate shallow waters (Bruce et al. 2010). Bottom type data derived from field notes (approximately 40 points per reef) were imported into ArcGIS (Environmental Systems Research Institute 2011) as point data and were subsequently used along with the bottom type data derived from the 2009 NOAA acoustic seabed mapping survey to supplement the delineation of bottom type contours of characteristic grain sizes (e.g., muddy sand). Bottom type information derived from the 2009 NOAA acoustic seabed mapping survey was utilized for the areas outside of the ACE constructed oyster reef polygons, while bottom type information from the field sampling notes taken by VIMS researchers was also utilized for the areas within the ACE constructed oyster reef polygons. This provided a baseline (i.e., pre-construction) bottom type dataset for integration into the HSI. This baseline bottom type dataset was necessary as some of the reefs within the sanctuary reef network were originally constructed on unsuitable bottom and have degraded since the 2009 NOAA acoustic seabed mapping survey.

Mean bottom salinity data were derived from a VIMS hydrodynamic model developed for tributaries of Chesapeake Bay (J. Shen, unpublished data). Salinity data were imported into ArcGIS (Environmental Systems Research Institute 2011) as point data (1 m nodes), and were subsequently converted to produce a raster grid (1 m x 1 m resolution). Bathymetric data were derived from a NOAA bathymetric digital elevation model of the Chesapeake Bay (1 m × 1 m resolution) (Bruce et al. 2010).

Within ArcGIS (Environmental Systems Research Institute 2011), each input variable layer was assigned suitability index values based on physiological tolerance ranges available in the literature, or from on-going research of the ACE reefs in the GWR. Data cells within each input variable layer were classified on a gradient between 0, indicating highly unsuitable habitat, and 1, indicating highly suitable habitat, with varying degrees of suitability in between. In order to ensure that the HSI would assign a particular location a value of ‘0’ (unsuitable) if any single input variable layer had a value of ‘0,’ the geometric mean of each input variable layer was calculated to produce the HSI. Mathematically, the HSI was computed as follows:

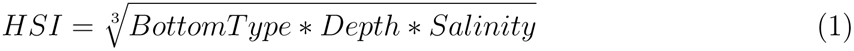

In Chesapeake Bay, a live adult oyster density of 50 m^−2^ is the abundance target for a restored reef to be considered successful by the Sustainable Fisheries Goal Implementation Team of the Chesapeake Bay Program (Sustainable Fisheries Goal Implementation Team 2011). Thus, live adult oyster density data derived from the 2011 VIMS monitoring survey of the ACE restored reefs in the GWR were used to validate the model.

The 2011 VIMS monitoring survey involved sampling HRR and LRR sanctuary oyster reefs (Figure 3) via patent tong. The sampling methodology followed standard procedures under Stratified Random Sampling (Cochran 1977, Thompson 2012). Reef types were then apportioned into strata (i.e., HRR, LRR) using information from side-scan sonar maps. Stratum area and variance estimates were used to generate random, stratum-specific nominal sampling sites and backup sites within a grid surrounding each of the reef polygons (Figure 3), using stratified random sampling with sample allocation proportional to stratum area and variance (Cochran 1977, Thompson 2012). Sampling sites were located by GPS coordinates, and sampled in the order in which they were generated to assure random sampling within each stratum. At each sampling site, the vessel was triple-anchored to maintain position. Next, a patent tong (1 m wide) was deployed, the sample was retrieved on a processing table aboard the vessel, and a photo taken of the sample with its identification number visible on a whiteboard. A complete 0.5 m^2^ section of the 1 m^2^ tong sample was rinsed and retained for lab processing. Samples were processed in the laboratory, rather than in the field, due to the high probability that individual oysters would not be easily seen in the field, resulting in biased (inaccurate) samples. Parameter estimates for density and abundance were obtained using the R statistics package (R Core Team 2015).

**Figure 3.**
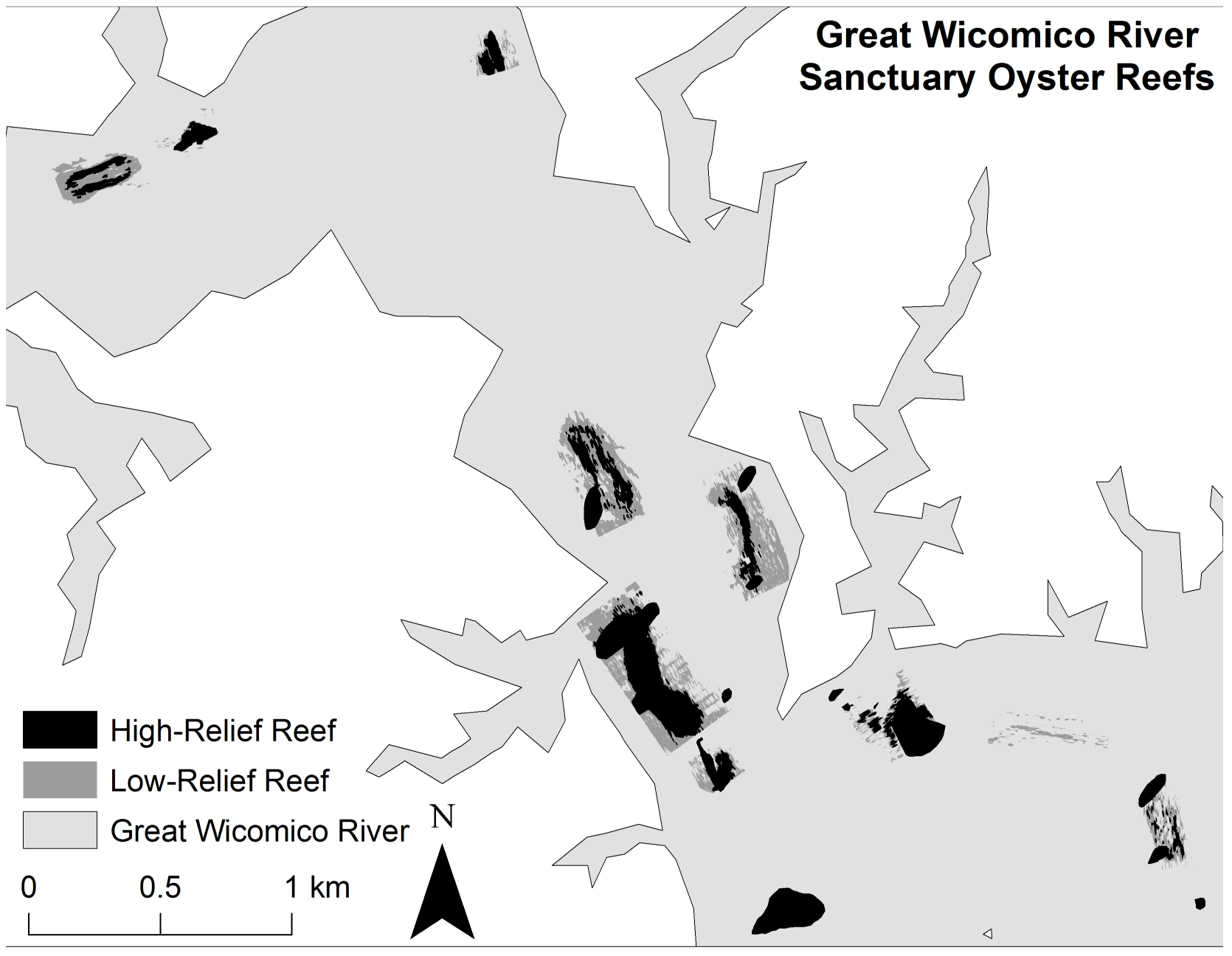
Distribution of high vertical relief reefs (HRR) and low vertical relief reefs (LRR) in the US Army Corps of Engineers network of sanctuary reefs in the Great Wicomico River.

## Results

### HSI distribution in the Great Wicomico River

Bottom type and depth were the primary drivers of a particular site’s suitability (Figure 4). The river’s bathymetry, which consists of a deep, soft bottom channel flanked by shallow, hard bottom shelfs mirrors the observed suitability trends. Despite the most suitable locations existing in shallow depth areas, patches of unsuitable bottom intermixed in these areas constrain their extent. Additionally, in upriver locations, lower salinity waters reduces the extent of highly suitable habitat.

**Figure 4.**
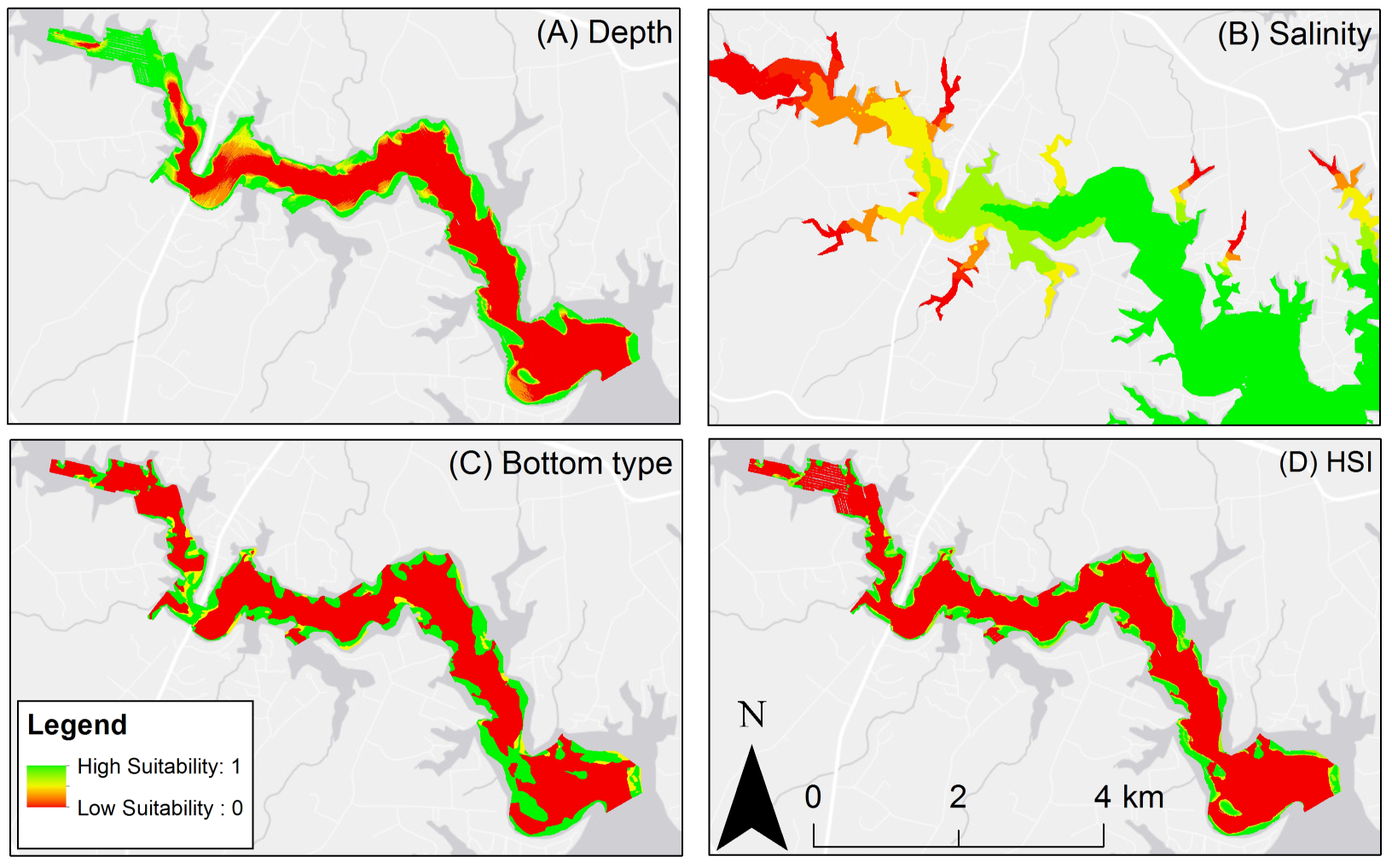
A) Depth suitability layer. B) Salinity suitability layer. C) Bottom type suitability layer. D) Habitat suitability index output, red areas are unsuitable oyster habitat, green areas are optimal oyster habitat.

### HSI calibration and validation

After computation of the HSI, the model was calibrated through linear rescaling of the HSI values to ensure the model ranged from 0 to 1. Live adult oyster density was a statistically-significant sigmoid function of the HSI (Figure 5), thereby validating the HSI as a robust predictor of suitable oyster reef habitat for rehabilitation or restoration in the GWR. In addition, high-relief reef had on average 103-116 more adults m^−2^ than low-relief reef at a given level of the HSI. For high-relief reefs, HSI values ≥0.3 exceeded the 50 live adult oysters m^−2^ target (Figure 5). For low-relief reefs, the HSI was generally able to predict live adult oyster densities that meet or exceed the target at HSI values ≥0.3. However, for low-relief reefs, there was more variation between the HSI value and the corresponding live adult oyster density than for high-relief reefs. To maximize the potential for success of oyster reef restoration and rehabilitation, suitable oyster habitat was defined as all areas with an HSI value ≥0.5, and marginal oyster habitat was defined as all areas with an HSI value ≥0.3.

**Figure 5.**
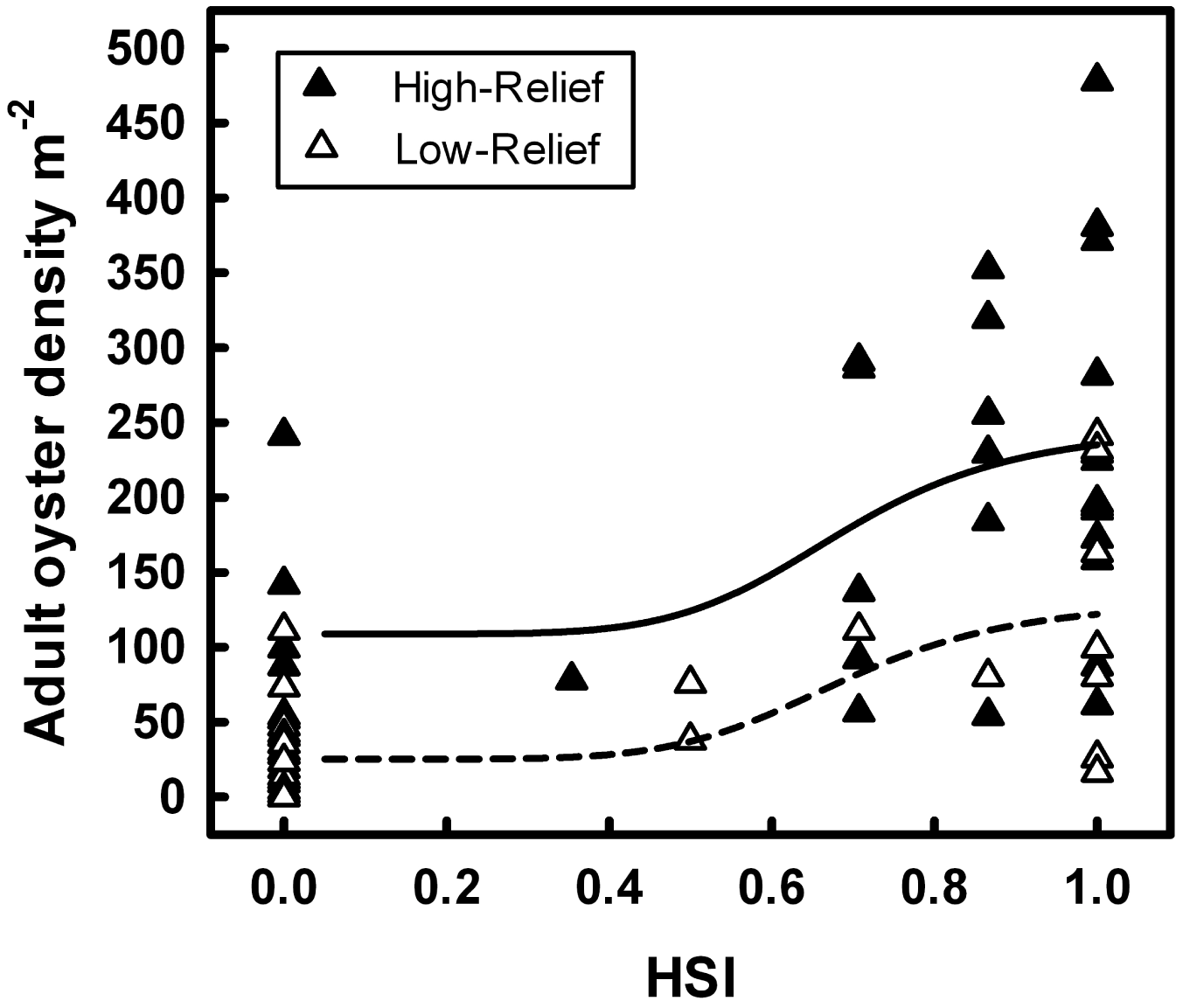
Sigmoid relationship between habitat suitability index score and associated live adult oyster density on HRR and LRR. The solid line represents the relationship for HRR and the dashed line represents the relationship for LRR.

### Suitable habitat

This study identified that approximately 157.8 ha of suitable oyster habitat (e.g., cells with a HSI value between 0.5 and 1.0) occur in the GWR. Approximately 2.4 ha of marginally suitable oyster habitat (e.g., cells with an HSI value between 0.3 and 0.5) occur in the GWR. Approximately 92.7 ha of suitable oyster habitat lay within private shellfish aquaculture leases, along with approximately 0.8 ha of marginally suitable oyster habitat. Within the 288.5 ha of public oyster harvest grounds, approximately 52.6 ha of suitable oyster habitat and 0 ha of marginally suitable oyster habitat exist. On the ACE high-relief reef, approximately 13.4 ha of suitable oyster habitat and approximately 0.2 ha of marginally suitable habitat exist. On the ACE low-relief reef, approximately 9.7 ha of suitable oyster habitat and approximately 0.2 ha of marginally suitable oyster habitat exist. It is important to note that, due to the lack of available bottom type data for shallow areas, the total amount of suitable habitat available in the GWR is greater than the amount estimated here. As the majority of shallow areas in the GWR are held in the form of private shellfish aquaculture leases, most of this additional suitable habitat is likely contained within the private shellfish aquaculture leases.

## Discussion

Habitat suitability indices provide a quantitative tool that integrates the best available environmental data and corresponding science to identify locations with the most potential for successful restoration (U. S. Fish and Wildlife Service 1981, Roloff and Kernohan 1999), but their efficacy depends critically on validation with independent data (Araújo and Guisan 2006, Brooks 1997, Tirpak et al. 2009, Reiley et al. 2014). Of the eight published HSI models used in native oyster restoration and fishery production (Table 1), none has been validated. Our key contributions are thus the (i) development, calibration and validation of a relatively simple HSI model for restoration of the eastern oyster *Crassostrea virginica,* (ii) demonstration of the value of including a key factor, in this case reef relief, in the implementation of the HSI model, and (iii) identification of oyster abundance as a sigmoid function of the HSI. The HSI was validated with data from a metapopulation undergoing restoration in the GWR (Schulte et al. 2009), and depended largely on bottom type, salinity, and water depth, a surrogate for dissolved oxygen concentration. Given the extensive effort being dedicated to eastern oyster restoration across the Atlantic and Gulf of Mexico coasts, our findings should enhance the efficiency and effectiveness of such efforts, and likely for other native oyster species worldwide.

Some of the previous oyster HSI models have included additional variables, such as turbidity and predation intensity (Table 2), for smaller or more intensively studied waterbodies, where these spatial data sets are available. For larger or less studied waterbodies, spatial data sets for large suites of environmental variables are often rare. Further, based on the wide range of environmental and biotic variables included in prior HSI models (Table 2), it is clear that a “one size fits all” approach to HSI modeling is inappropriate. While many variables that determine habitat suitability overlap between species and systems, variables that are significant drivers in a given system may not be as critical as other variables. For instance, of the four oyster HSI models developed for specific systems (i.e., Caloosahatchee Estuary, Florida by Barnes et al. (2007), Hudson River, New York by Starke et al. (2011), Mission-Aransas Estuary, Texas by Pollack et al. (2012), Mississippi River, Louisiana by Soniat et al. (2013)), only a single variable was used by all four (i.e., average salinity), yet they used a total of seven other variables that were not in common (Table 2).

The eastern oyster provides an excellent example of this variability in requirements in that its geographic range encompasses the (i) Atlantic coast from Canada to Florida, (ii) Gulf of Mexico coast, and (iii) Caribbean from the Yucatan Peninsula to the West Indies (Buroker 1983), which spans no less than five biogeographic provinces containing a diverse suite of habitats across temperate, subtropical and tropical areas (Kennedy et al. 1996). When using HSI models that incorporate a small subset of variables, such as only salinity and substrate (e.g., Soniat et al. (2013), Swannack et al. (2014)), it is especially valuable to conduct model validation. Such simple models trade incorporation of large suites of environmental layers (for which there are rarely comprehensive spatial datasets for large areas) for broader spatial coverage, with the underlying assumption being that oyster habitat suitability can be adequately described by salinity and substrate alone. In the case of Chesapeake Bay, for which seasonal anoxia and hypoxia are prevalent in deeper waters, omission of a variable that encapsulated dissolved oxygen (i.e., water depth) may lead to an HSI erroneously overstating the extent of suitable oyster habitat for restoration. In this case, water depth was an effective, if imperfect, surrogate for dissolved oxygen, and was largely accountable for the performance of the HSI model along with bottom type. Bathymetric information is often available for most waterbodies, and an understanding of the relationship between depth and dissolved oxygen concentrations can be useful to eliminate areas of hypoxia or anoxia in restoration efforts by use of HSI models. In the case of the GWR, seasonal hypoxia in areas deeper than 4 m, reduced salinity in upriver locations, and subsidence of reef material in areas of soft sediments had previously been identified as priority factors that could negatively impact the success of oyster restoration. Thus, developing an HSI that integrates the variables known to be major drivers of restoration success in a particular system with subsequent validation and model refinement is the optimal, robust approach.

Given the commonly stated goal for oyster restoration projects of oyster abundance and biomass enhancement, live adult oyster density data derived from the 2011 survey of the ACE restored reefs in the GWR were used to validate the model. Our use of live adult oyster density data from seven years post-restoration allowed us to avoid biases resulting from anomalous recruitment events. In addition, when using oyster density data to conduct HSI model validation, it is important to differentiate live adult oyster density and total live oyster density. Immediately following restoration, oyster sanctuaries can experience major recruitment pulses that can temporarily inflate total oyster density with size structure skewed towards high densities of recruits and sub-adults, which have reduced probabilities of survival relative to adults (Puckett and Eggleston 2012).

Our division of the analysis of live adult oyster densities and corresponding HSI values into low‐ and high-relief reef categories was necessary as some of the low-relief reefs constructed in the GWR had degraded due to sedimentation and subsidence into mud seabottom. For low-relief reefs, reef persistence depends greatly on whether live oyster shell accretion outpaces subsidence and sedimentation (Schulte et al. 2009, Jordan-Cooley et al. 2011, Colden and Lipcius 2015, Lipcius et al. 2015). In several cases, low-relief reefs in the GWR with HSI values ≥0.3 exhibited low live adult oyster densities. These low-relief reefs containing suitable oyster habitat should be examined as candidates for rehabilitation to high-relief reefs through the placement of additional shell material on the low relief sites. The majority of high-relief reefs in the GWR have persisted due to the accretion of additional shell material, which allows the reef matrix to outpace subsidence and sedimentation and thereby maintain a positive shell budget (Powell et al. 2012). These finding lends support to the existence of bistability (i.e., alternative stable states-persistent reef or degraded reef-existing under similar environmental conditions) on the restored oyster reefs in the GWR. In the persistent state, high-relief reefs (i) offset heavy sedimentation, (ii) promote larval settlement and survival, juvenile survival and growth, and adult disease resistance and survival, (iii) lead to elevated adult abundance and reef accretion, and (iv) have high adult abundance which enhances settlement and recruitment, thereby constituting a positive feedback cycle. In contrast, in the low-relief degraded state (i) there is heavy sediment deposition, (ii) thick sediments on the reef inhibit larval settlement and juvenile/adult survival, (iii) low adult abundance limits reef accretion, and (iv) depressed adult abundance and reef height preclude high settlement and facilitate heavy sediment accumulation, leading to a downward spiral towards a degraded reef and local extinction of the oyster population (Jordan-Cooley et al. 2011).

Live adult oyster density was a sigmoid function of the HSI for both low‐ and high-relief reefs. The sigmoid relationship differs from the linear relationship between density and HSI in previous validated studies (Reiley et al. 2014, Cho et al. 2012). The sigmoid relationship implies, for both low‐ and high-relief reef categories, a stasis in oyster density below an HSI of 0.3, and a rapid increase in oyster density with increasing HSI to a value likely corresponding with a site’s carrying capacity. Low-relief reefs in areas with an HSI ≤0.3 are unlikely to exceed performance standards, such as that of the Sustainable Fisheries Goal Implementation Team (2011)-live adult oyster density target of 50 m^−2^. However, in areas of low-relief reef with an HSI ≥0.3, density rapidly increases to nearly double the 50 live adult oysters m^−2^ target in low-relief reef ares with an HSI of 1. Contrasting this, for high-relief reef in areas with an HSI as low as 0, the 50 live adult oysters m^−2^ target is exceeded with a rapid increase in density to nearly 5x the target at an HSI of 1, although the long-term sustainability of reefs in areas with low associated HSI values is questionable. This finding also provides further support to the existence of bistability on the restored oyster reefs in the GWR in that under similar environmental conditions (i.e., for a given level of HSI), low‐ and high-relief reefs remain differentiated by oyster density.

With the increasing availability of spatial datasets for environmental variables in marine and estuarine systems, habitat suitability indices will likely continue to be developed to inform species conservation and management. However, as cautioned by the USFWS shortly after the development of the first HSIs in the 1980s, the performance of these models must be quantitatively assessed via an independent dataset. This study provides a robust framework for HSI model development and validation, which can be refined and applied to other systems and previously developed HSIs to improve the efficacy of native oyster restoration.

## Acknowledgments

We thank the VIMS Marine Conservation Biology and Community Ecology Labs for their assistance with oyster sampling. We thank KW Theuerkauf and anonymous reviewers for helpful comments that improved this manuscript, and David Bruce of the National Oceanic and Atmospheric Administration, Chesapeake Bay Office for technical assistance with the benthic habitat maps. Funding for the project and completion of the manuscript were provided by the College of William and Mary Roy R. Charles Center for Academic Excellence through an Honors Fellowship, James Monroe Scholarship, and Center for Geospatial Analysis Student Research Grant awarded to SJT, as well as grants from the U.S. Army Corps of Engineers, Norfolk District, the National Science Foundation’s Mathematical Biology program, and the National Oceanic and Atmospheric Administration, Chesapeake Bay Office to RNL. This is Contribution No. XXXX of the Virginia Institute of Marine Science, College of William & Mary.

